# Region-specific regulation of glucocorticoid and mineralocorticoid receptor signaling in a mouse model of oral contraceptive exposure

**DOI:** 10.64898/2026.06.15.731933

**Authors:** Kristen M. Schuh, Mackenzie G. Woock, Megan J. Vaandrager, Elizabeth G. Romano, Yutong He, Daniella Ludmir, Natalie C. Tronson

**Affiliations:** Psychology Department, University of Michigan, Ann Arbor MI 48109

**Author notes:** Corresponding author: Natalie C. Tronson, Address: Psychology Department, University of Michigan, 530 Church st, Ann Arbor, MI 48109. **Funding:** University of Michigan Eisenberg Family Depression Center Oscar Stern Award to NCT; University of Michigan LSA-OVPR funds to NCT; F31 HD114532 to KMS.

## Abstract

Combined oral contraceptives (OCs), containing synthetic estrogen and a progestin such as levonorgestrel (LVNG), are widely used, and up to 10% of users experience adverse mood states and increased depression risk. It is well-established that OCs modulate the hypothalamic-pituitary-adrenal (HPA) axis and blunt the cortisol responses to acute stress. This interaction with stress regulatory pathways is one mechanism by which OCs might impact mood. Here, we used a mouse model of OC exposure (ethinyl estradiol (EE) + LVNG) to investigate how OCs affect regulation of the diurnal CORT cycle and stress-related signaling in the dorsal and ventral hippocampus and paraventricular nucleus of the hypothalamus (PVN). We found that EE+LVNG did not alter basal corticosterone (CORT) levels, but impaired glucocorticoid receptor (GR) - mediated negative feedback in the dexamethasone suppression test. Molecular analyses revealed distinct, region-specific effects. In the dorsal hippocampus, EE+LVNG enhanced glucocorticoid receptor (GR)-dependent gene signaling and prolonged *Fkbp5* induction. In the ventral hippocampus, EE+LVNG enhanced mineralocorticoid receptor (MR)-dependent signaling and reduced stress-induced corticotropin-releasing factor expression. In the PVN, EE+LVNG reduced *MR* expression and modulated MR-dependent signaling. Together, these findings demonstrate that chronic OC exposure disrupts GR- and MR-dependent regulation across stress-related brain regions and impairs glucocorticoid feedback, providing potential mechanisms by which OCs blunt stress responsivity, modify long-term HPA-axis function, and increase susceptibility or resilience to stress and depression.

---

Hormonal contraceptives, including combined oral contraceptives (OCs), are the most prescribed drugs worldwide, and are used by more than 400 million women [1]. Up to 10% of users experience adverse mood states and increased risk for depression, yet despite more than 60 years of OC use, we still do not how OCs modify, or who is susceptible to OC-mediated increases in risk for depression.

Combined OCs are typically comprised of ethinyl estradiol (EE), and a synthetic progestin, which varies by formulation, and primarily suppress hypothalamic–pituitary–gonadal axis and ovulation. OCs also regulate the hypothalamic–pituitary–adrenal (HPA) axis, which controls response to stress. OC users exhibit blunted cortisol responses to acute stress [2–7], an effect that evident in rodent models of OCs [8–10]. Given the role of stress in the pathogenesis of depression [11], regulation of HPA axis is a plausible mechanism by which OCs modify depression risk.

In the brain, the paraventricular nucleus of the hypothalamus (PVN) initiates the stress response [12], and in response to glucocorticoid receptor (GR) response to CORT, the dorsal and ventral hippocampus (dHPC, vHPC) inhibit the PVN, thereby suppressing ongoing HPA output [13,14]. dHPC, vHPC, and PVN are thus strong candidate loci for OC-modulation of stress [15].

Cortisol and corticosterone (CORT) signal though mineralocorticoid receptor (MR), a high affinity receptor engaged under basal conditions, and GR, a lower-affinity which is activated during stress [16]. Coordinated MR and GR signaling in PVN [17] and HPC [18–20] controls initiation and resolution of the stress response, and disruptions can impair negative feedback, contributing to prolonged CORT production, and increased vulnerability to stress-related psychopathology [21].

Transcripts downstream of MR and GR are indices and regulators of receptor activity (Figure 1). *Period* 1 (*Per1)* and *Serum/glucocorticoid-regulated kinase 1* (*Sgk1)* are primarily GR targets [22,23]; whereas *Jun dimerization protein 2* (*Jdp2)* is MR-dependent [23]. *FK506-binding protein 51* (*fkbp5*) is downstream of both MR and GR, and functionally binds to GR, inhibiting GR sensitivity [24]. Finally, corticotropin releasing factor (*crf*) and 11β-hydroxysteroid dehydrogenase type 1 (*hsd11β1*) regulate feedforward aspects of the stress response [25].

**Figure 1.**
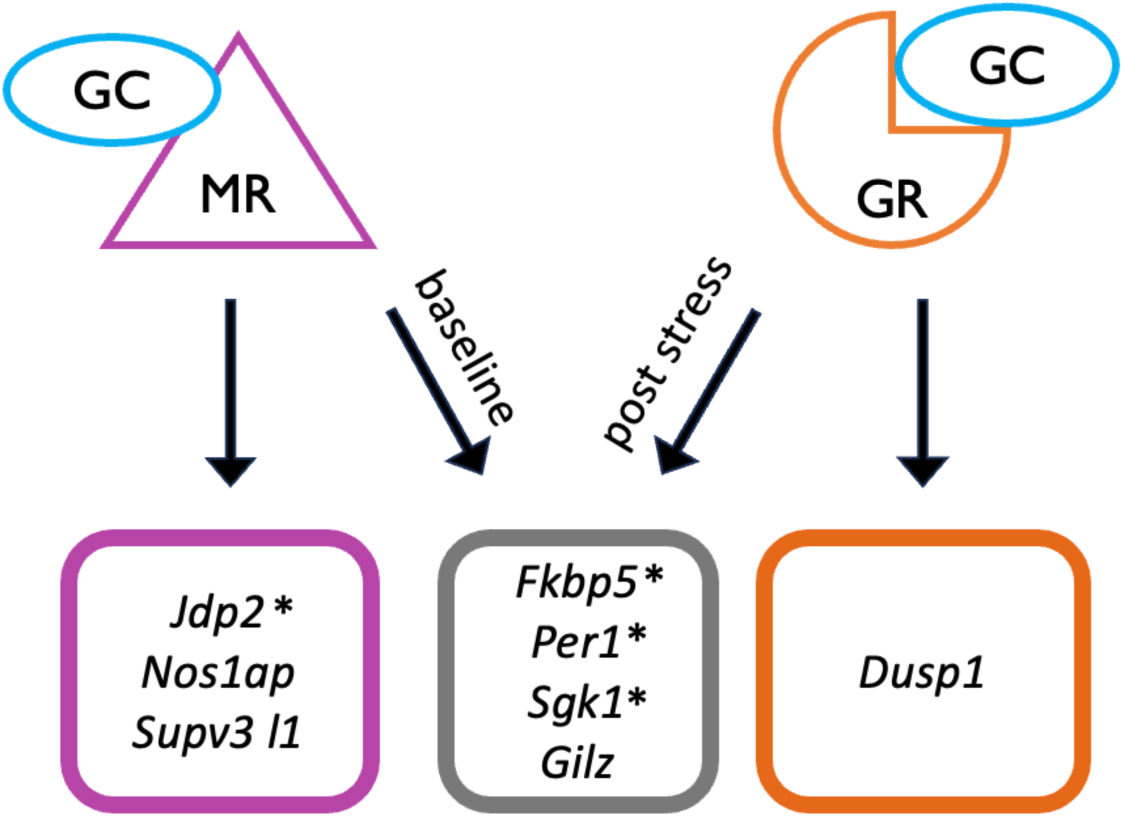
MR- and GR-dependent downstream transcriptional targets. *Measured in this manuscript. GC: glucocorticoids, MR: mineralocorticoid receptor, GR: glucocorticoid receptor.

FKBP5 is of particular interest: it plays a central role in regulation of GR-signaling, and because circulating FKBP5 is increased in OC users [3], suggesting a mechanism by which OC exposure regulates stress. Polymorphisms in FKBP5 are a known risk for stress-related disorders [26–29], and is thus a potential mechanism linking OCs, stress dysregulation, and depression risk. In the brain, the precise role of *fkbp5* expression is region-specific. In the hippocampus, *fkbp5* expression is tightly coupled to MR activity and when elevated, reduces GR sensitivity [30], potentially decreasing HPC suppression of PVN and resulting in persistent CORT levels after stress. Conversely, in the PVN, *Fkbp5* enhances the CORT response to stress response [31].

Importantly, the critical role of brain circuits in HPA axis regulation [12,13,32] makes identifying the mechanisms by which OCs interact with stress a difficult question to study in humans [9,33–35]. Thus, in this paper, we use a mouse model of OC exposure [8], one of a growing number of rodent models developed to address the central effects of hormonal contraceptives [36–39]. We evaluated GR-mediated negative feedback using the dexamethasone (DEX) suppression test (DST) and quantified OC-induced changes in *Fkbp5*, *Nr3c1* (GR), *Nr3c2* (MR), *Per1*, *Sgk1*, *Jdp2*, *Crf*, and *Hsd11b1* expression in the PVN, dHPC, and vHPC. Based on prior findings [3,5,8], we hypothesized that OC exposure increases *Fkbp5* expression, enhances MR signaling, and decreases GR activation in the dHPC and vHPC; while in the PVN, OCs will decrease *Fkbp5* and *Crf* expression after stress.

## METHODS

### Animals

We used adult female C57Bl6/N mice (9-10 weeks on arrival) from Envigo (Indianapolis, IN). Mice were individually housed in the colony room with access to standard chow and water *ad libitum* and acclimatize for 7 days. The colony room, adjacent to behavioral testing rooms, was maintained at 20 ± 2 °C with a 12h 0700: 1900 light/dark cycle (lights on at 0700 h). All protocols and treatments were approved by the University of Michigan Institutional Animal Care and Use Committee.

### Oral Contraceptive Hormone Treatment

OC treatment: 250μL of 10% sucrose with a suspension of ethinyl estradiol (EE, 0.01875μg/mouse, 0.9375μg/kg per 20g mouse)) and levonorgestrel (LVNG, 0.75μg/ mouse, or ∼37.5μg/kg per 20g mouse) [8].

Control treatment: 250μL of 10% sucrose.

Treatments were delivered daily in small cell culture dishes placed in home cages prior to lights out (1700-1900h). Mice were observed to confirm that all mice consumed the solution (within minutes). Daily OC treatment continued throughout all experimental procedures.

### Diurnal CORT Rhythmicity and Dexamethasone Suppression Test (DST): Experiment design

Mice (n=20) received EE+LVNG/control for 2 weeks prior to jugular vein and abdominal cannulation surgeries, with 5 days to recover and habituate to Culex automatic blood collection chambers. Hourly blood samples (24 samples, 25μL per sample) were collected on day 1. On day 2, dexamethasone (DEX, 40 ug/kg) was administered through i.p. cannulae at 10 am, and blood was sampled at 5’, 15’, 30’, 2 hours, and then every 4 hours post-DEX injection. Serum samples (35/animal) were stored at -80°C. The UM Mouse Metabolic Phenotyping Center (MMPC) performed this experiment.

### OC modulation of GR- and MR-dependent gene expression: Experiment design

Mice received EE+LVNG or sucrose control for at least 4 weeks prior to 60 mins of stress. In one experiment mice (N=48) were exposed to restraint stress with dHPC, vHPC, and PVN dissected 30mins after the end of restraint. In a second experiment, mice (N=48) were exposed to either restraint stress or footshock stress and dHPC, vHPC, and PVN dissected 4 hours after the end of stress. In a third experiment, mice (N=16) were exposed restraint stress, brains perfused 60min post stress, and PVN gene expression assessed using RNAScope.

### Acute Stress

#### Acute Restraint Stress

Mice were briefly anesthetized isoflurane drop box, and taped prone to a flat surface for one hour [40].

#### Acute Footshock Stress

Mice were placed into a chamber (MedAssociates) for 60 min, and received 13 pseudorandom 1mA, 1-3sec footshocks (adapted from [41]).

### Blood and tissue collection

Blood was allowed to clot at room temperature for 30 min, then centrifuged at 15294 g (12000 RPM) for 15 minutes, serum supernatant extracted and stored at -20°C.

For qPCR, mice were rapidly decapitated, dHPC and vHPC were dissected, and PVN was punched out, flash frozen and stored in the -80°C freezer.

For RNAScope, one hour after the end of the acute restraint stress, mice were deeply anesthetized (250 mg/kg of Avertin), transcardially perfused with 4% paraformaldehyde. Brains were post-fixed in 4% paraformaldehyde for 24 hours, and stored at 4°C.

### ELISA

CORT and ACTH ELISAs (Arbor Assays #K014-H and K072-H) were used following the manufacturer’s protocol [8].

### Quantitative real-time PCR

qPCR was conducted as previously described [42]. mRNA was extracted from homogenized tissue (PureLink RNA Mini Kit; Cat. No. 12183020; Invitrogen, Carlsbad, CA), quantified, and purity assessed (A260/280 > 1.80) using UV spectrometry (BioSpectrometer Basic; Eppendorf, Hamburg, Germany). 800ng of cDNA was synthesized for v/dHPC, and 240ng for PVN (QuantiTect Reverse Transcriptase Kit; Cat. No. 205314; Qiagen, Hilden, Germany).

Power SYBR Green master mix (Cat. No. 4368702; Applied Biosystems, Foster City, CA) and 10uL reactions (QuantStudio™5 System; Cat. No. A28140; Applied Biosystems) were used.

Primer sequences: Table 1.

**Table 1.**
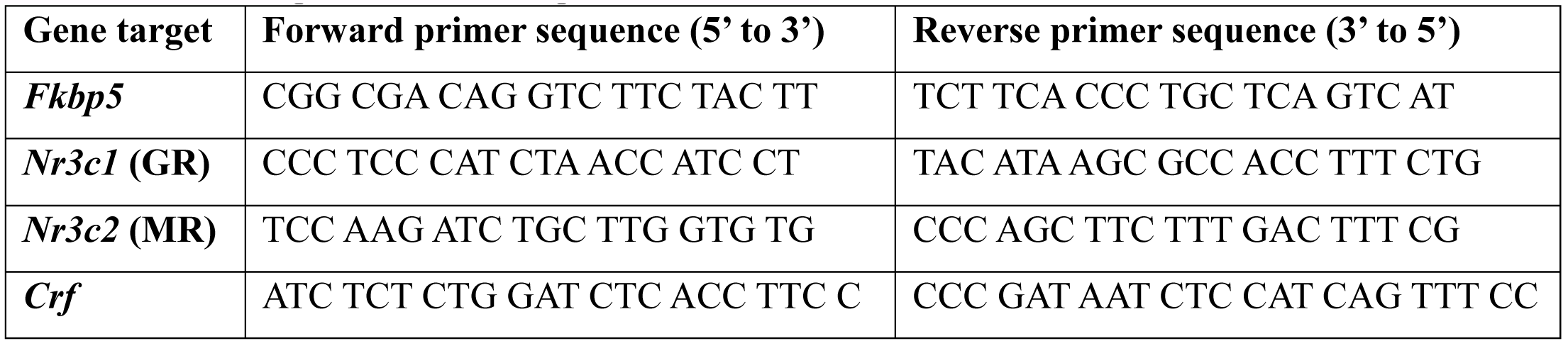
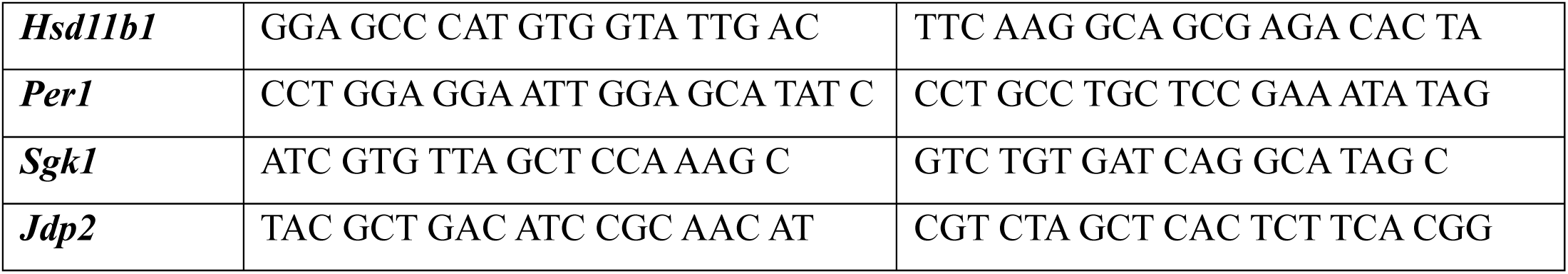
Primer sequences used for qPCR.

### RNAscope

10 µm coronal sections were mounted onto slides and stored at -80°C. RNAscope fluorescent *in situ* hybridization (Multiplex Reagent Kit (cat. no. 320850, Advanced Cell Diagnostics, Newark, CA) was conducted according to the manufacturer’s protocol [43] to quantify cellular mRNA expression for GR (probe mm-Nr3c1), MR (probe mm-Nr3c2-C2) and *Fkbp5* (probe: mm-Fkbp5-C3). Sections were counterstained with DAPI. Confocal imaging and ImageJ was to analyze expression in the PVN [44]

### Statistical Analysis

GraphPad Prism 8 and SPSS (v.29) were used for statistical analyses. ANOVA, repeated measures ANOVA, and MANOVA were used throughout, with post-hoc Bonferroni tests to control for multiple tests.

## RESULTS

### EE+LVNG does not significantly alter circadian basal corticosterone levels

Basal CORT levels did not differ between EE+LVNG-treated and control mice (Figure 2A (Time: F(3.54, 53.39) = 17.28, p<0.0001; OC: F(1, 16) = 0.154, p=0.70; time x OC: F(12, 181) = 1.86, p<0.05). This interaction was driven primarily by the 3:00am CORT measurement, a timepoint which coincides with routine cleaning activity in adjacent rooms and was consistent across all cohorts over a 3-month period. At this time point, control mice showed higher CORT levels relative to EE+LVNG-treated mice, a difference that approached significance (p=0.059). We further compared the increase of CORT concentrations from the 1:00am (baseline) to 3:00am peak (Figure 2B: Time, F(1, 31) = 34.62, p<0.0001; OC treatment (p=0.05); Time x OC interaction: F(1, 31) = 4.88, p<0.05). This replicates our previous finding [8] that EE+LVNG blunts the CORT response to stress.

**Figure 2.**
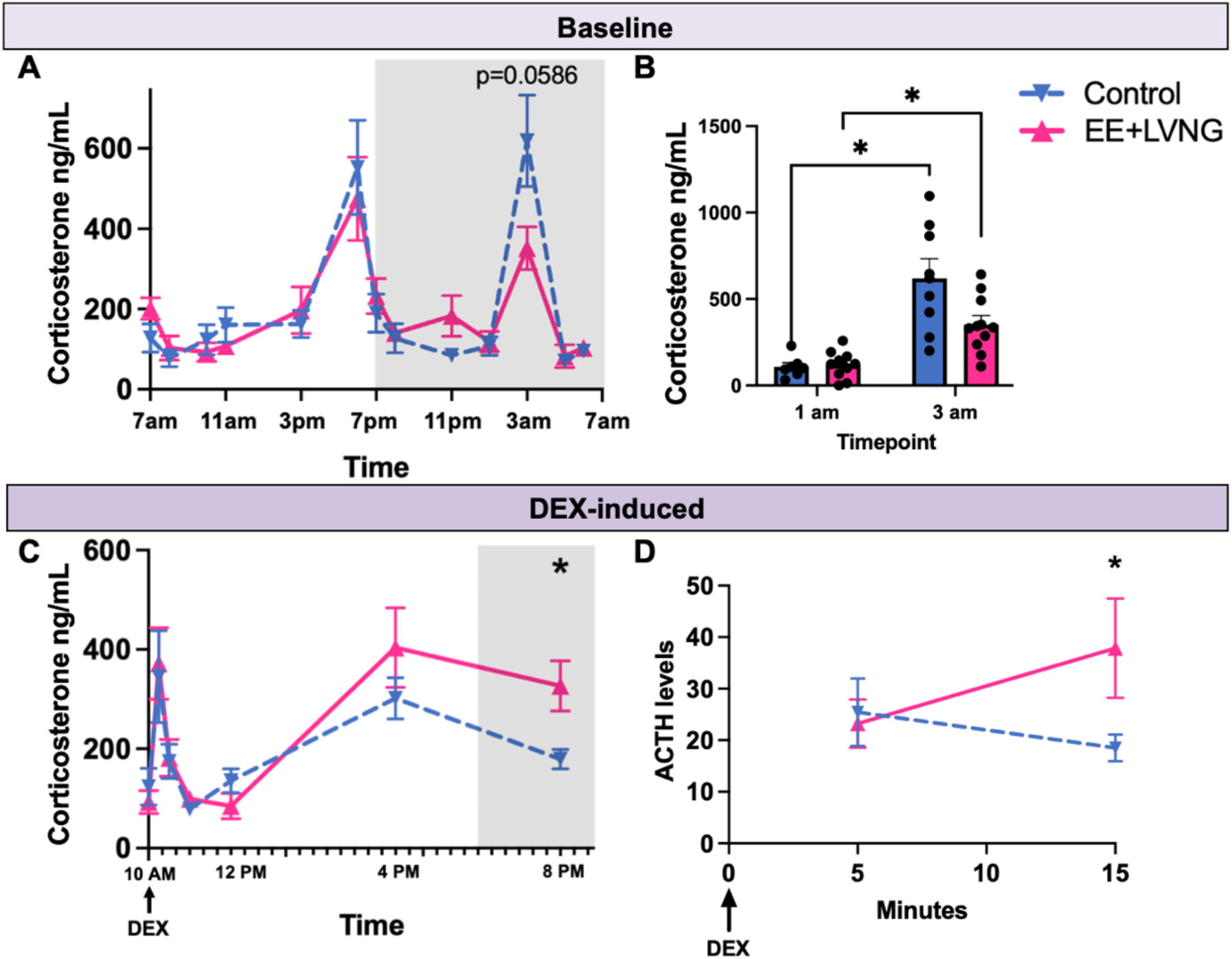
Maintained circadian CORT rhythm with OCs, but impaired CORT suppression in the DST (A) Circadian CORT levels (ng/mL) over 24 hours. (B) OC-treated mice showed a blunted CORT response at 3:00AM due to an environmental stressor. (C) OC-treated mice displayed significantly higher CORT concentrations at 8:00 pm (p<0.05), indicating impaired CORT suppression. (D) 15-min after DEX, EE+LVNG mice exhibited higher ACTH levels vs controls. Darker bars represent dark phase. *p<0.05.

### EE+LVNG reduced dexamethasone suppression

Both treatment groups showed an initial rise and fall of CORT following DEX administration. A significant main effect of time was observed, (F(3.021, 43.30) = 12.26, p<0.001). There was no effect of OC treatment (F(1,15)<1), nor a time x OC treatment interaction (F(6, 86) = 1.17, p=0.33). Notably, EE+LVNG-treated mice exhibited reduced suppression at later time points, with higher CORT concentrations compared to controls at 8:00 pm (p<0.05) (Figure 2C). These findings suggest that EE+LVNG exposure impairs glucocorticoid negative feedback, consistent with reduced sensitivity to DEX.

EE+LVNG-treated mice exhibited significantly elevated plasma ACTH levels at 15-min post-injection compared to controls (p<0.05; Figure 2D), whereas no group differences were observed at the 5-min timepoint.

### Dorsal hippocampus: EE+LVNG enhanced stress-induced GR activity

30 min post-stress: A multivariate analysis of variance (MANOVA) revealed a significant main effect of stress (Pillai’s Trace = 0.77, F(9, 30) = 11.36, p<0.0001, η_p_^2^ = 0.77), indicating robust early transcriptional activation. There was no significant multivariate main effect of OC treatment nor an interaction between stress and OC treatment. Univariate analyses showed that stress increased *Fkbp5* expression (F(1, 39) = 5.472, p<0.05, η_p_^2^ = 0.11), an effect primarily driven by EE+LVNG-treated mice (p<0.01; Figure 3B). Similarly, stress enhanced GR-dependent downstream signaling, as evidenced by elevated *Per1* expression (F(1, 39) = 10.61, p<0.01, η_p_^2^ = 0.21) again driven by EE+LVNG mice (p<0.01; Figure 3Ci). However, *Nr3c1* (GR) expression was not affected by stress or EE+LVNG treatment. GR-dependent gene target, *Sgk1*, was elevated by stress (F(1,38) = 26.14, p<0.0001, η_p_^2^ = 0.41), but not EE+LVNG treatment (p=0.80). MR expression and its downstream gene target, *Jdp2*, were not affected by stress nor OC treatment (Figure 3Ai, 3D). Stress increased MR:GR ratio (F(1,40) = 10.09, p<0.01, η_p_^2^ = 0.18), an effect potentially driven by EE+LVNG-treated mice (p<0.01). However, the lack of difference in control mice is likely due to a single data point. Neither *Crf* nor *Hsd11b1* was affected by stress or OC treatment (Figure 3Ei-ii).

**Figure 3.**
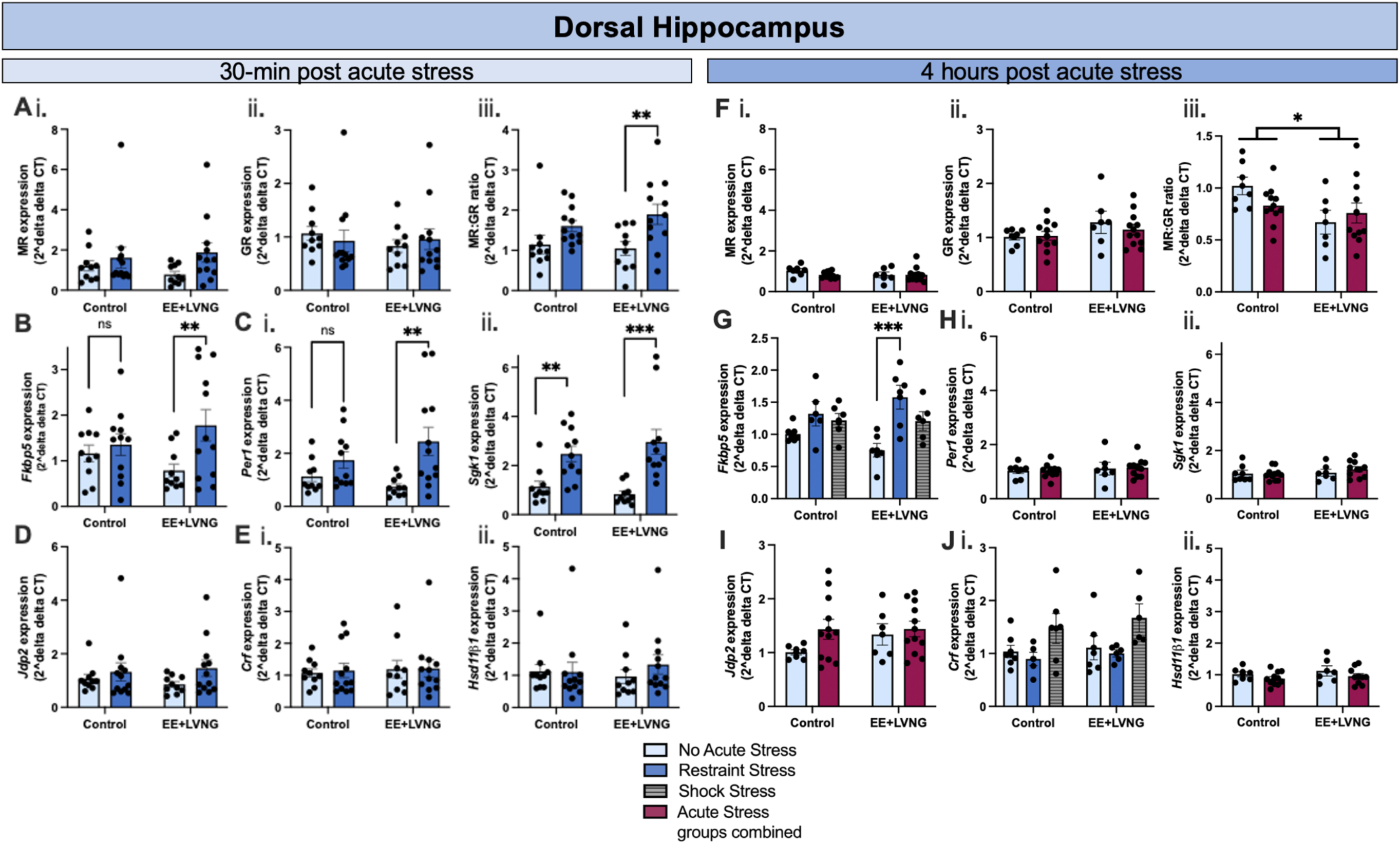
OC*s* modif*y* stress-induced gene expression in dorsal hippocampus. (A-E) **30 minutes post stress:** (A) (i) MR (i), GR, and (iii) stress increased MR:GR ratio, an effect driven by EE+LVNG-treated mice. (B) Fkbp5 expression was increased by stress, primarily in EE+LVNG-treated mice. (C) GR-dependent downstream targets (i) Per1 and (ii) Sgk1 were upregulated by stress, with greater induction of Per1 in EE+LVNG mice. (D) Jdp2 and (E) (i) Crf and (ii) Hsd11b1 were not significantly different across groups. (F-J) **4 hours post stress:** (F) (i) MR, (ii) GR, (iii) OC-exposure reduced the MR:GR ratio. (G) Fkbp5 remained elevated in stressed mice, particularly in OC and restraint-stressed mice. (H) Per1 and (I) Sgk1 expression returned to baseline. (J) (i) Jdp2 was unchanged, (ii) Crf was increased by stress, (iii) Hsd11b1 showed no significant 4 hours after stress. Error Bars ± SEM. **p<0.01, ***p<0.001

4 hours post-stress: MANOVAs revealed a significant main effect of stress when restraint and shock groups were combined (Pillai’s Trace = 0.67, F(9, 22) = 5.06, p<0.001, η_p_^2^ = 0.67), and for each stressor independently (Pillai’s Trace = 0.95, F(18, 42) = 2.10, p<0.05, η_p_^2^ = 0.47). Univariate analyses showed that neither EE+LVNG nor stress modulated MR or GR expression (Figure 3Fi-ii); however, EE+LVNG exposure decreased MR:GR ratio (F(1, 31) = 5.42, p<0.05, η_p_^2^ = 0.14; Figure 3F(iii)). *Fkbp5* expression remained elevated (main effect of stress, F(2, 29) = 9.624, p<0.001, η_p_^2^ = 0.37), particularly in EE+LVNG restraint-stressed mice (p<0.001; Figure 3G). GR-dependent downstream signaling (*Per1* and *Sgk1*) had returned to baseline (Figure 3Hi-ii), indicating that GR-specific activity is transient, whereas *Fkbp5*, downstream of both MR and GR (Figure 1), remains elevated for longer. Neither *Jdp2* nor *Hsd11b1* expression were affected by stress or OC exposure (Figure 3I, 3Jii). *Crf* expression was increased by stress (F(2, 28) = 5.198, p<0.05, η_p_^2^ = 0.38; Figure 3Ji). These findings demonstrate that the dorsal hippocampus mounts both rapid and sustained transcriptional responses to stress, with EE+LVNG exposure amplifying these effects.

### Ventral hippocampus: EE+LVNG enhanced stress-induced MR activity

30min post-stress: A MANOVA revealed a significant main effect of stress (Pillai’s Trace = 0.88, F(9, 19) = 15.57, p<0.001, η_p_^2^ = 0.88). There was no significant multivariate main effect of OC treatment (p=0.28) nor a stress x OC interaction (p=0.09). Univariate analyses showed no modulation of MR expression (Figure 4Ai); however, MR-dependent gene target (*Jdp2*) was elevated by stress (F(1, 39) = 7.755, p<0.01, η_p_^2^ = 0.17), an effect driven by EE+LVNG mice (p<0.01; Figure 4D). At baseline, *GR* expression was reduced by EE+LVNG exposure (p<0.05), and there was a significant stress x OC interaction (F(1, 35) = 4.235, p<0.05, η_p_^2^ = 0.11): stress increased GR expression only in OC mice (p<0.05; Figure 4Aii). GR-dependent downstream signaling was also increased by stress (*Per1*: F(1, 38) = 23.93, p<0.0001, η_p_^2^ = 0.34 and *Sgk1*: F(1, 39) = 53.79, p<0.0001, η_p_^2^ = 0.63; Figure 4Ci-ii). Neither *Fkbp5* expression nor MR:GR ratio was changed (Figure 4B; 4Aiii).

**Figure 4.**
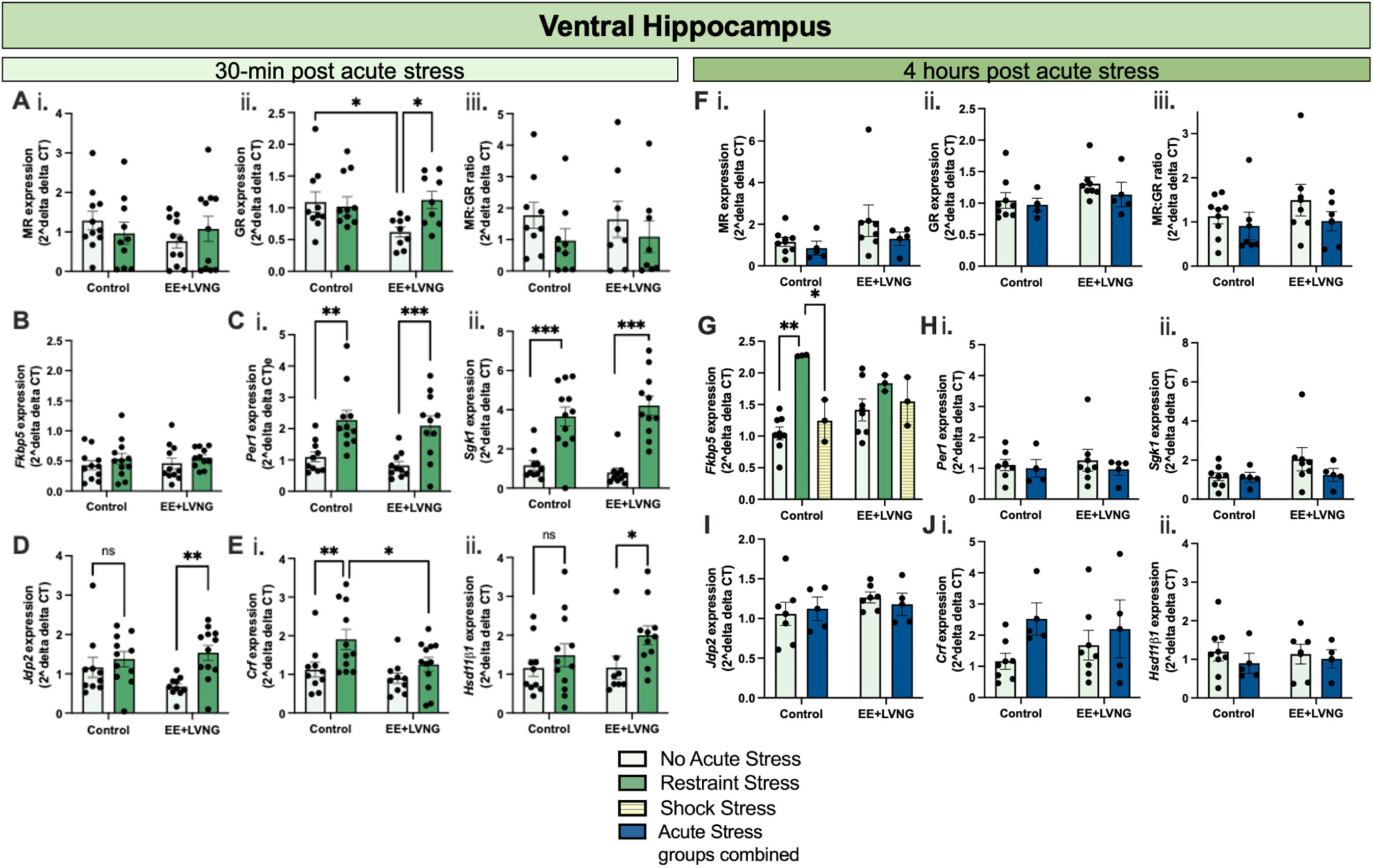
OCs modify stress-induced gene expression in ventral hippocampus. (A-E) **30 mins post-stress:** (A) MR (i), GR (ii), and MR:GR ratio (iii). EE+LVNG exposure decreased GR expression at baseline. Stress increased GR expression, with effects driven by EE+LVNG-treated mice. (B) *Fkbp5* expression was unchanged. (C) GR-dependent downstream targets (i) *Per1* (ii) *Sgk1* were robustly upregulated by stress. For (D) *Jdp2,* (E) (i) *Crf* and (ii) *Hsd11b1: Jdp2* and *Hsd11b1* were elevated by stress, an effect driven by OC-treated mice. *Crf* was attenuated by OC exposure. (F-J) **4 hours post-stress:** (F) MR (i), GR (ii), and MR:GR ratio (iii) were unaffected by stress and EE+LVNG. (G) *Fkbp5* expression was elevated in stressed mice, an effect driven by control restraint-stressed mice. (H) *Per1* (i) and *Sgk1* (ii) returned to baseline. (I) *Jdp2*, (J) *Crf* (i), and *Hsd11b1* (ii) returned to baseline. Error bars ± SEM. *p<0.05, **p<0.01, ***p<0.001

*Crf* expression was elevated by stress (F(1, 39) = 8.074, p<0.01, η_p_^2^ = 0.18), and OC treatment reduced *Crf* expression (F(1, 39) = 4.667, p<0.05, η_p_^2^ = 0.18; Figure 4Ei). Stress increased *Hsd11b1* expression (F(1, 37) = 4.61, p<0.05, η_p_^2^ = 0.13; Figure 4Eii), an effect driven by the EE+LVNG group (p<0.05).

4 hours post-stress: MANOVA results did not reveal a significant main effect of stress when restraint and shock groups were combined (p=0.11); however, when the stress groups were separated, there was a main effect of stress (Pillai’s Trace = 1.59, F(18, 12) = 2.55, p=0.05, η_p_^2^ = 0.79). We could not replicate the baseline *GR* differences, and the stress x OC interaction was no longer present (Figure 4Fii). MR- and GR-dependent downstream targets had generally returned to baseline levels (Figure 4H-I). However, stress increased *Fkbp5* expression at this time point, this time driven by control restraint-stressed mice (F(2, 17) = 7.692, p<0.05, η_p_^2^ = 0.57; Figure 4G). *Crf* and *Hsd11b1* had returned to baseline at this time point (Figure 4Ji-ii). These findings indicate that the ventral hippocampus undergoes early MR- and GR-dependent transcriptional activation, with EE+LVNG exposure amplifying MR-dependent effects.

### PVN: EE+LVNG modulated MR activity

In the PVN, we analyzed transcriptional changes using RNAscope 1 hour after acute restraint stress. We observed a significant main effect of OC treatment on *Nr3c2* (MR) expression, F (1, 12) = 9.829, p<0.01, with EE+LVNG-treated mice showing reduced MR expression relative to controls (Figure 5Aii). Post-hoc analysis confirmed that control mice exposed to stress exhibited higher MR expression compared to EE+LVNG-stressed mice (p<0.01). In contrast, neither *Fkbp5* nor *Nr3c1* (GR) expression differed across groups (Figure 5Aiii, 3.5Av), and MR:GR ratio was unaffected (Figure 5Aiv).

**Figure 5.**
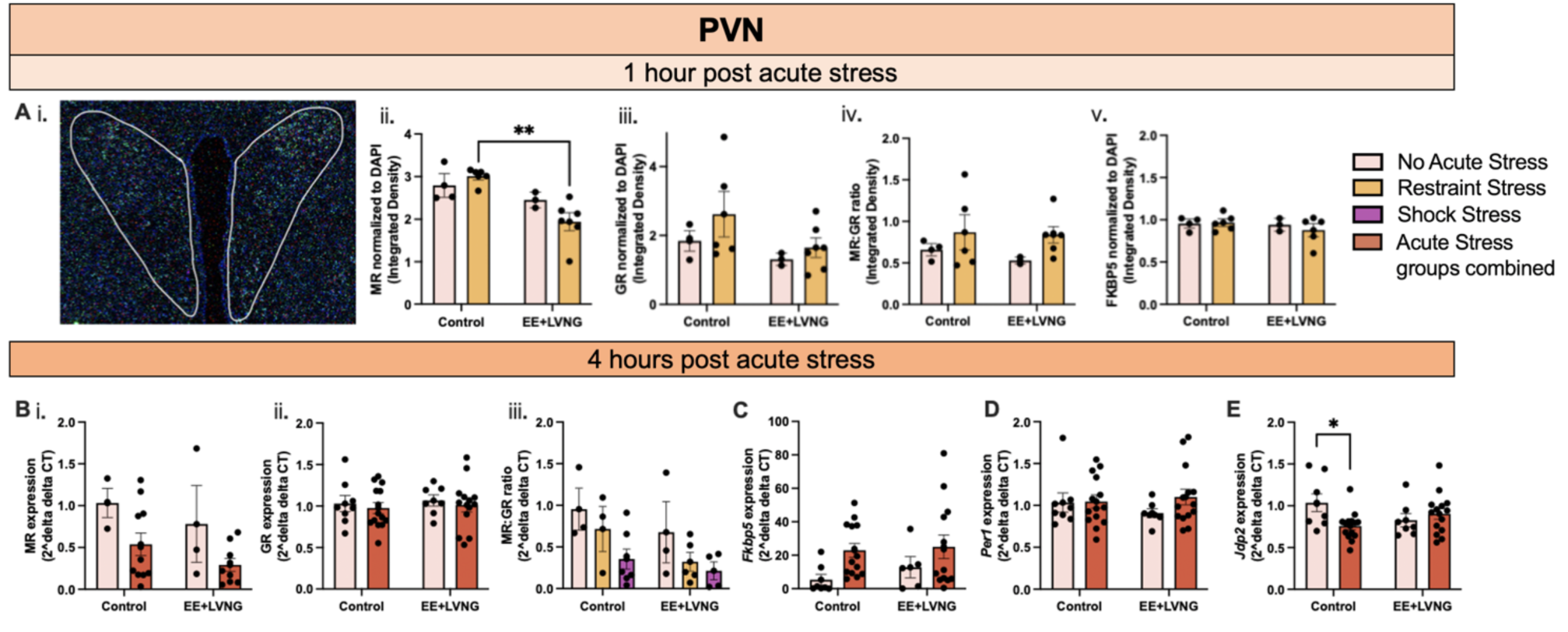
OC*s* modif*y* stress-induced gene expression in PVN. (A) Gene expression (integrated density) normalized to DAPI for MR, GR, Fkbp5, and MR:GR one hour after acute restraint stress. (i) Representative PVN RNAscope image. (ii) MR expression was decreased by EE+LVNG, (iii) GR, (iv) MR:GR ratio, and (v) Fkbp5 expression were not *different across condition*. red=GR, cyan= Fkbp5, green=MR, Dark blue= DAPI. (B-E) 4 hours after stress (qPCR) (B) (i) MR expression was attenuated. (ii) GR expression; (iii) MR:GR ratio was decreased. (C) Fkbp5 increased; (D) Per1 expression (E) Jdp2 decreased in control, but not EE+LVNG-treated mice, Error Bars ± SEM. *p<0.05. **p<0.01

4 hours after acute restraint or shock stress, MANOVA on qPCR results showed no main effect of stress when restraint and shock groups were combined (p=0.81); however, in the model with the stress groups separated, a main effect of stress emerged (Pillai’s Trace = 0.92, F(12, 34) = 2.43, p<0.05, η_p_^2^ = 0.46). The stress x OC interaction approached significance (Pillai’s Trace = 0.82, F(12, 34) = 1.97, p=0.06, η_p_^2^ = 0.41). Univariate analyses showed that stress decreased MR (F(1, 22) = 6.284, p<0.05, η_p_^2^ = 0.22; Figure 5Bi) and increased *Fkbp5* (F(1, 39) = 7.755, p<0.01, η_p_^2^ = 0.13; Figure 5C) expression. Neither stress nor OC exposure modified GR expression (Figure 5Bii), or GR-dependent transcription (*Per1*; Figure 5C). Stress decreased MR:GR ratio (F(2, 19) = 3.856, p<0.05, η_p_^2^ = 0.25; Figure 5Biii). There was a significant stress x OC interaction for Jdp2 expression (F(1, 39) = 7.755, p<0.01, η_p_^2^ = 0.13; Figure 5E). In control mice only, stress decreased MR-dependent downstream signaling (p<0.05).

## DISCUSSION

We used our mouse OC model [8] to examine how OC affects diurnal CORT regulation, GR-mediated negative feedback, and to identify molecular mechanisms by which OCs regulate the response to acute stress. One important finding is that chronic EE+LVNG exposure impairs negative feedback regulation, indicated by incomplete DEX suppression and elevated ACTH levels following treatment. Distinct, region-specific transcriptional changes mirror this effect: EE+LVNG initially enhanced stress-induced GR-dependent signaling and subsequently increased the GR-regulatory co-chaperone *fkbp5* in the dHPC; enhanced stress-induced MR-dependent signaling in the vHPC, and disrupted MR expression and MR-dependent signaling in the PVN.

EE+LVNG did not broadly alter basal CORT or the cyclicity across 24 hours (Figure 2A-B). Nevertheless, across five independent cohorts, we observed a rise in CORT at approximately 3:00 a.m., coinciding with the time when cleaning staff were active in adjacent rooms. The blunting of CORT in OC-treated mice is consistent with our prior data [8]. These data further suggest that EE+LNG differentially impacts mechanisms underlying diurnal regulation and stress–related signaling [8,9].

EE+LVNG treatment impaired overnight DEX suppression, (Figure 2C), paralleling reports in some OC users exhibiting hypercortisolism-like symptoms [45] and supporting our hypothesis that OCs decrease GR sensitivity and negative feedback. Elevated ACTH after DEX administration (Figure 2D), suggest that OCs may also increase pituitary release of ACTH. Together, these findings indicate that OCs do not simply blunt the stress response but may fundamentally alter the set point of negative feedback regulation of the HPA axis, increasing susceptibility to stress dysregulation in some individuals.

Consistent with the decreased GR function observed the DST, and with elevated FKBP5 in OC users [3], we observed early enhancement of the GR-mediated gene expression, followed by persistent increases only in *Fkbp5* expression, suggesting that after the initial response to stress, GR sensitivity decreases, resulting in decreased inhibition of PVN, and persistently elevated CORT in the hours after an stressor.

In the vHPC, OCs promoted MR activity and the local CORT signaling via *Hsd11b1*.

Enhanced MR activity together with reduced *Crf* expression is consistent with blunted systemic CORT response to stress.

In PVN, OC exposure also predominantly modified MR-related transcription, albeit in a more complex manner than vHPC. Four hours, but not one hour after stress, EE+LVNG prevented the stress-triggered decrease in *Jdp2* expression in control animals. Given the PVN’s central role in initiating ACTH release and coordinating neuroendocrine output [12], changes in MR activity may contribute to increased hypothalamic sensitivity to modulatory input from upstream pathways, including hippocampal inhibition. These transcriptional changes in the PVN demonstrate that EE+LVNG exposure modifies HPA axis feedback regulation, a potential mechanism by which EE+LVNG blunts the CORT response to stress.

The regulation of MR in PVN and vHPC may be due to a LVNG-specific effect.

Endogenous progesterone acts as an MR antagonist, whereas LVNG does not [9]. Thus, by suppressing endogenous progesterone and replacing it with a non-antagonistic progestin, EE+LVNG may effectively impact MR signaling in vHPC.

We also observed differential effects of stress and OCs on dHPC and vHPC. The vHPC is speculated to be more directly engaged in emotional regulation and inhibitory control of the HPA axis [14,46,47], whereas the dorsal hippocampus may respond more rapidly to stressors and support cognitive aspects of stress processing [48,49]. Nevertheless, dHPC is more often studied for its role in HPA-axis regulation, and few studies have compared the stress-regulatory functions of these subregions. Our data and that of others, suggest that MR activation may exert opposing effects in dHPC versus vHPC [50].

*Jdp2* induction in the vHPC and attenuation in the PVN following stress indicate that MR signaling in this region is not restricted to basal conditions, challenging the classical view that MR solely governs basal glucocorticoid tone while GR mediates stress responses[29,51]. Recent work suggests that the majority of hippocampal MR are dynamically regulated by both circadian and stress-related glucocorticoid fluctuations [52,53], indicating that MR participates in both basal and stress-induced signaling. Thus, the differential engagement of MR versus GR signaling across brain regions likely reflects region-specific receptor composition and regulation rather than simple CORT availability.

One critical future direction is identifying individual difference factors contributing to risk for adverse outcomes from OCs[54]. Our data support the idea that Fkbp5 may be a key mechanism for stress-regulation during OC use [3]. Thus, *Fkbp5* polymorphisms known to increase risk for stress-related disorders may also increase risk for OC-triggered depression.

We demonstrated that EE+LVNG exposure disrupted glucocorticoid feedback regulation at in the periphery, characterized by impaired DEX suppression, and elevated ACTH; and in the brain, shown by dHPC-specific shifts in GR- and vHPC/PVN specific changes in MR-dependent gene expression. Together, these findings suggest that OCs recalibrate the feedback architecture of the HPA axis, that may contribute to susceptibility to depression in some individuals, while increasing resilience in others.

## Acknowledgements

University of Michigan Animal Phenotyping Core (MMPC), who conducted the Culex diurnal CORT and DST blood collection study. Drs. Hanna Carmon and Caitlin Posillico for qPCR expertise and training. Dr. Shigeki Iwase and the Human Genetics Department for equipment usage. Dr. Dave Bridges for assistance and advice throughout.

Finally, our lab manager Amy (Hoi Kei) Choi who assisted in everything, every day.

## Author Contributions

KMS designed and conducted experiments including ELISA, qPCR, RNAScope, collaborated with MMPC, conducted statistical analyses, wrote the manuscript; MGW, MV, EGR, YH, DL assisted with all experiments, specifically MGW: qPCR and associated statistical analyses; MV, DL: ELISA analyses; EGR, YH, DL qPCR. NCT designed experiments, conducted statistical analyses, and co-wrote manuscript.

